# Unraveling Why We Sleep: Quantitative Analysis Reveals Abrupt Transition from Neural Reorganization to Repair in Early Development

**DOI:** 10.1101/827212

**Authors:** Junyu Cao, Alexander B. Herman, Geoffrey B. West, Gina Poe, Van M. Savage

## Abstract

Sleep serves disparate functions, most notably neural repair, metabolite clearance and circuit reorganization, yet the relative importance of these functions remains hotly debated. Here, we create a novel mechanistic framework for understanding and predicting how sleep changes during ontogeny (why babies sleep twice as long as adults) and across phylogeny (why mice sleep roughly five times that of whales). We use this theory to quantitatively distinguish between sleep used for neural reorganization versus repair. We conduct a comprehensive, quantitative analysis of human sleep using total sleep time, cerebral metabolic rate, brain size, synaptic density, and REM sleep (used here to also refer to Active Sleep in infants and children). Our findings reveal an abrupt transition, between 2 and 3 years of age in humans. Specifically, our results show that differences in sleep across phylogeny and during late ontogeny (after 2 or 3 years in humans) are primarily due to sleep functioning for repair or clearance, while changes in sleep during early ontogeny (before 2 - 3 years in humans) primarily support neural reorganization and learning. Moreover, our analysis shows that neuroplastic reorganization occurs primarily in REM sleep but not in NREM. In accordance with the developmental role of neuroplasticity, the percent of time spent in REM sleep is independent of brain size across species but decreases dramatically as brain size grows through development. Furthermore, the ratio of NREM sleep time to awake time emerges as a new invariant across development. This developmental transition and fundamental shift across ontogeny and phylogeny suggests a complex interplay between developmental and evolutionary constraints on sleep.

## Introduction

The pervasiveness of sleep during development and throughout the animal kingdom suggests it is a biological process that is necessary for survival. Although we spend approximately a third of our life asleep, its explicit physiological and evolutionary function remains unclear with myriad hypotheses having been postulated (1–28). Two of the leading hypotheses are that sleep enables (i) the repair and clearance needed to correct and prevent neuronal damage (1–12, 29–32) and (ii) the neural reorganization necessary for learning and synaptic homeostasis (13–21). These hypotheses are compelling because neither of these processes can be easily achieved in waking states and there is supporting empirical evidence that they occur during sleep.

For instance, prolonged sleep deprivation can lead to death in rats (25), dogs (26), fruit flies (27), and even humans (28). These extreme cases are believed to result from damage to neuronal cells caused by metabolic processes that are not avoided or remedied because clearance of damaging agents and repair occur primarily during sleep (6,29). Moreover, a recent hypothesis related to neuronal damage from metabolic processes is that sleep drives metabolic clearance from the brain (7). Because the brain lacks a penetrating lymphatic system, cerebrospinal fluid recirculates through the brain and removes interstitial proteins (8, 9), likely through meningeal lymphatics (33). Furthermore, the concentration of *β*−amyloid(*Aβ*) is higher in the awake state than during sleep, suggesting that wakefulness is associated with producing (*Aβ*) (10, 11) while sleep is associated with its clearance. This view is further supported by the 60% increase in interstitial space associated with sleep (7).

There is also substantial and direct evidence that sleep promotes neuroplastic reorganization (15) related to learning and consolidating memory and also regulates synaptic re-scaling. For instance, neuron firing sequences that encode spatial maps learned during awake periods are replayed during sleep (17, 34–38). In addition, sleep facilitates the growth of learning-associated synapses and the homeostatic weakening and pruning of seldom-used synapses (6, 21). More generally, two recent studies demonstrate that sleep regulates the cycling of proteins related to synaptic functioning (39, 40).

Comparative, developmental, physiological, and human studies have all been fruitfully used to address questions about the nature of sleep (1–28) and each has given different and often complementary insights into its function (41). However, because data are seldom analyzed in a way that connects them with mathematical models or quantitative predictions, conclusions about the function of sleep have remained slow to evolve. In this context, we develop a general theory for the function of sleep that provides a framework for addressing several fundamental questions, such as: What are the relative roles of repair and reorganization during sleep, and do these change during ontogenetic development?

An important quantitative observation is that sleep times systematically decrease with body mass across mammals (42, 43). Moreover, the fraction of time spent in REM sleep (also referred to as active sleep) does not change with brain or body mass (42). Since increasing body mass strongly correlates with decreasing mass-specific metabolic rate (i.e., metabolic rate per unit mass) and therefore to a decreasing rate of cellular damage, this strongly suggests that less sleep time is needed for repair and maintenance in larger animals. These empirical observations led two of us (42) to develop a quantitative mechanistic theory for understanding the origins and function of sleep across species based on the central role played by metabolism in both damage and repair. This work suggested novel analyses of the empirical data on brain size and brain metabolic rate, both of which depend non-linearly on body size, and showed that both brain size and brain metabolic rate are better predictors of sleep time than body size. This provided strong evidence that sleep is primarily associated with repair of the brain rather than with the other parts of the body. Specifically, we predicted that the ratio of sleep time to awake time should decrease with brain size as a power law whose exponent is 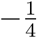, and consequently that it should decrease with body weight with an exponent of 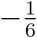, both in good agreement with data. The scaling exponent of 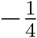 for brain size corresponds to the same scaling as mass-specific metabolic rate in the brain. The theory also provides a quantitative understanding for why the proportion of REM sleep does not change with either brain or body mass.

The major focus of this paper is to address the intriguing question as to whether these general relationships for sleep times remain valid during growth, implying that ontogeny recapitulates phylogeny, or whether new patterns emerge during development reflecting a different dynamical origin for sleep? More pointedly, during both development and across species, sleep times systematically decrease with brain and body size. But do they do so at similar rates, and are they attributable to the same underlying dynamics? If new patterns emerge, what do those patterns reveal about neurological development and the growth of the brain? To answer these questions, we derive a quantitative ontogenetic sleep model across species that combines *both* ontogeny and phylogeny in a single framework, and use this model to guide the analysis of human sleep data from birth to adult. We compare our new findings with previous empirical and theoretical results for how sleep changes across phylogeny to ask if explanations for why a mouse sleeps roughly five times more than a whale can also be used to explain why babies sleep roughly twice as long as adults.

Although previous studies focus on total sleep time, and how it is partitioned between REM and NREM, and how these change during growth (44, 45), we are unaware of any systematic quantitative mechanistic models for how or why these change as children grow. Here, we combine the most comprehensive published data on sleep throughout human development and across species with a new mechanistic model to elucidate the function of sleep, reveal how it dramatically changes during early growth, and show how this is related to brain development.

In the following section, we develop a framework for modeling neural repair/metabolite clearance and reorganization during sleep, and show how brain metabolic rate depends on the number of synaptic connections between neurons. Moreover, we propose a general quantitative model for how sleep time changes with brain mass ontogenetically. Next, we describe the sources and collection of our data as well as the statistical and numerical methods for how the data were analyzed. We collate and integrate data for total sleep time, REM sleep time, brain weight, body weight, cerebral metabolic rate, and synaptic density based on a systematic review of the literature. The resulting dataset spans from 0-15 years of age and cumulatively represents about 400 data points. We then use the empirical data to discover patterns of sleep during ontogeny, compare them with phylogenetic patterns, and test predictions from our theoretical framework. In so doing, we:

1. develop distinct quantitative theories for both neural repair/clearance and neural reorganization,
2. use extensive human sleep and brain data from birth to adult to cleanly test and discriminate among theories,
3. provide strong evidence of a remarkably sharp transition in the purpose of sleep at about 2.4 years of age from the purpose of sleep being primarily for neural reorganization that occurs during the Active Sleep/REM cycle in early development to sleep being primarily for neural repair and metabolite clearance in late development.

Finally, we explain our conclusions and discuss remaining questions and future directions.

## 1 Framework for predicting sleep times and testing sleep functions

Our conceptual, quantitative framework for how sleep changes as brains increase in size and age through development is grounded in key hypotheses about the dominant function of sleep being for neural repair/clearance and/or reorganization. We explain simple mathematical equivalencies that lead to specific, baseline predictions for scaling exponents that encapsulate how ratios of REM, NREM, and total sleep times change with brain size.

### 1.1 Theory of sleep for neural repair

We previously constructed a mathematical theory that focused on neural repair in adult brains and empirically tested a suite of predictions for how characteristic times for sleep change with brain and body size across species (42). The theory, which we first briefly review, has been supported by investigations in recent years (7). It has long been postulated, and there is increasing empirical and theoretical evidence favoring it, that neural repair or clearance of metabolic wastes is an important function of sleep (7, 46). One theory, for instance, suggests that sleep plays the role of regulating oxidative stress in the brain by restoring and repairing neurons damaged by this oxidative stress (47). It has also been found that the production of oxidating agents in the brain during awake time promotes sleep (48–51). Recently, in the zebrafish it was found that chromatin movement, supporting remodeling and repair, happens only during sleep (29).

The basis of our theory is that the total amount of damage incurred and/or the accumulation of damaging agents during wakefulness must be reversed or counteracted by repair during sleep. Unlike other organs, it is crucial for the continuing functionality of the entire organism that neurological damage be faithfully repaired. The total damage that is generated during awake time is proportional to the mass-specific metabolic rate of the brain, *B*_*b*_, (effectively, the average metabolic rate of a cell) multiplied by the total time awake, *t*_*A*_. To counteract this, the total amount of repair or clearance accomplished during sleep is the power density, *P*_*R*_, allocated to repair or clearance during sleep multiplied by the total brain volume *V*_*b*_(∝ *M*_*b*_) and total sleep time, *t*_*S*_. Assuming that nearly all damage must be repaired or cleared in order for the brain to continue to function normally and with high fidelity throughout growth and adulthood, the total damage or accumulation of damaging agents must be balanced by the total repair or clearance.

This leads to

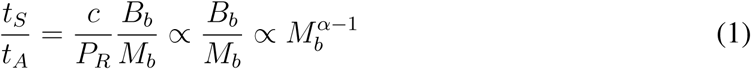

where *c* is a constant that incorporates the efficiency of repair processes together with the fraction of metabolic rate that leads to damage via free radicals, metabolic waste, or vessel damage. *P*_*R*_ is a local, cellular quantity and is assumed to be independent of body or brain size. Consequently, the predicted scaling exponent for sleep times is completely determined by the scaling of brain metabolic rate, 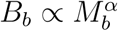 and therefore by its scaling exponent, *α*. For simplicity, we have here also assumed that all damage or accumulation of damaging agents occurs during wakefulness and that all repair and clearance occurs during sleep. This theory can straightforwardly be generalized to include damage during sleep and thereby to show that the dominant scaling relationships do not change. Indeed, this leads to an estimate that damage rates during sleep are about 1*/*3 of those during wakefulness (42).

Eq. (1) predicts that the *ratio* of sleep time to awake time follows a simple power-law relationship, which is well supported below by data. Indeed, the theory predicts that sleep time, *t*_*S*_, by itself does not obey pure power-law behavior with respect to brain size. Rather, it is the *ratios* of sleep to awake times or REM times that do (Supp. Info. 1). Because of this functional form, traditional plots in the literature for either *t*_*S*_ or ln(*t*_*S*_) versus ln(*M*_*b*_) are predicted to have much greater variance than for corresponding ratios and, more importantly, to be much more difficult to interpret.

Another key prediction of this theory based on neural repair is the invariance of the fraction of REM sleep

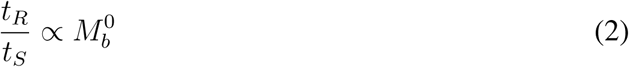

This pattern strongly holds across species. (42). Consequently, testing whether it remains valid during development will help reveal whether sleep during growth is primarily driven by neural repair or by some other function such as neural reorganization.

### 1.2 Theory of sleep for neural reorganization

During early development the brain is undergoing extensive changes in size, architecture, and cellular makeup. One of the major changes is in synaptic plasticity, which is greatest during early development after which it declines to a baseline adult level (52). This corresponds to neural reorganization, somato-cortical pathway development, and pruning that underlie the experience-dependent plasticity and learning necessary for adult behavior (53). Sleep is required for the consolidation and optimization of learning and governs underlying synaptic processes including synapse formation, sizing, and pruning (20, 21).

We now develop the theory for those aspects of neural reorganization related to sleep by focusing on the fundamental need to process *information*. Analogous to the theory for repair, the basis of the theory is an accounting and balancing of the rate of information being sensed by the body with the rate at which it is being processed by the brain. A key component of this model is that the amount of information needed to be processed is sensed through the entire body because stimuli are received from all parts of the body via pain, heat, cold, pressure, etc. On the other hand, the number of inputs that can be processed is constrained by the brain and its metabolic rate, *B*_*b*_, because the central nervous system is where the neural reorganization is occurring and where the information encountered by the peripheral nervous system is incorporated through synaptic structures for memory and learning. This crucial insight that information input is associated with the entire body, whereas its processing is localized in the brain, leads to a mismatch in the scaling of all of the sleep processes because brain size and brain metabolic rate scale non-linearly and differently with both body size and whole body metabolic rate, *B* (Supp. Info. 2 and (41)).

A core question is whether synaptic plasticity and information processing are occurring during just one or both of the two main stages of sleep, NREM and REM. Studies show that both are important for learning and memory, although their relative roles remain a topic of intense debate (54–58). Evidence suggests that REM may be more associated with local circuit changes reflecting memory consolidation (59) while during NREM global synaptic homeostasis and inter-region memory transfer may dominate (20, 60). Other evidence (58, 61, 62) suggests that synaptic pruning and reconnection primarily take place during REM sleep, while other results and arguments have posited that NREM sleep is when pruning and reorganizing occur (56).

Given this controversy, we derive separate predictions assuming either the primacy of REM or NREM sleep for neural reorganization. Consequently, our theory provides a quantitative test and a means for distinguishing between these two opposing hypotheses for the importance of REM versus NREM sleep by analyzing developmental sleep data. For simplicity, we present our equations in terms of REM sleep, since the NREM predictions are obtained by simply swapping NREM for REM everywhere in the following equations.

Assuming (i) that local neural reorganization associated with changes in synaptic density primarily occurs during REM sleep, (ii) that idealized synaptogenesis occurs uniformly across the brain, and (iii) that information exchange is directly tied to energy use, we relate the amount of information sensed by the body during wakefulness when the organism is being exposed to myriad stimuli to the amount being processed by the brain during sleep.

Defining Δ*E*_*I*→*σ*_ as the energy needed to convert a unit of information acquired by sensory systems to synaptic changes in the brain, and *f*_*I*_ as the fraction of the total metabolic rate required for sensing that information, then information is being transmitted to the brain at a rate given by (*f*_*I*_*B*)*/*Δ*E*_*I*→*σ*_. Consequently, the total amount of information generated while awake is proportional to (*f*_*I*_*Bt*_*A*_)*/*Δ*E*_*I*→*σ*_.

This information has to be processed during sleep by synapses (63). On average, each synapse processes information at a rate *ν* that, like all processes directly linked to brain metabolism, is expected to scale inversely with its mass-specific metabolic rate *B*_*b*_*/M*_*b*_. That is, inversely with cellular metabolic rate (42). Assuming first that local synaptic changes occur during REM sleep, *t*_*R*_, the total information processed is (*N*_*σ*_*t*_*R*_*ν*) ∝ (*N*_*σ*_*B*_*b*_*t*_*R*_*/M*_*b*_), where *N*_*σ*_ is the total number of synapses in the brain. We neglect terms related to synapses being formed and pruned within that same sleep-wake cycle because this number will be very small over such a relatively short time scale. Finally, equating the information processed during sleep with information sensed while awake and rearranging terms, we have

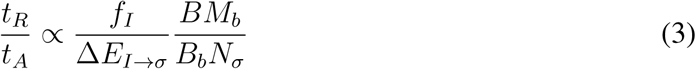

To determine how this ratio scales with brain and body size, we first recognize that the parameters *f*_*I*_ and Δ*E*_*I*→*σ*_ represent energies and percentages that are typically invariant with respect to size, in contrast to the scaling of biological rates and times (64–67). To express our result in terms of brain size, *M*_*b*_, we note that across species (42) and throughout development (Fig. S1) brain size scales nonlinearly with body size as approximately *M*_*b*_ ∝ *M* ^3*/*4^, so combining this with the canonical allometric relationship for whole body metabolic rate, *B* ∝ *M* ^3*/*4^ (valid through ontogeny), gives *B* ∝ *M*_*b*_. In the following section, we further argue that *N*_*σ*_ ∝ *B*_*b*_, thereby predicting the scaling of the ratio of REM sleep time to awake time:

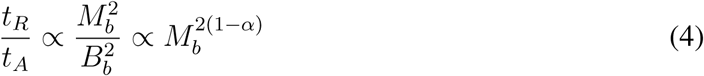

where *α* is the scaling exponent that relates brain metabolic rate to brain size. This can be re-expressed in terms of the ratio of REM sleep time to total sleep time (which is invariant across species (42)):

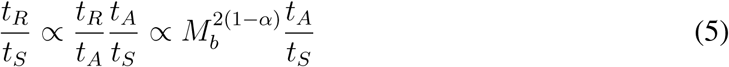

Thus, an empirical determination of how the ratio *t*_*A*_*/t*_*S*_ scales with brain size provides a prediction for the fraction of time spent in REM sleep across development. As we shall show below, these relationships provide a good description of the data and an important test of the theory. Furthermore, by simply switching NREM sleep time, *t*_*NR*_, with REM sleep time, *t*_*R*_, in the above equations, we also have the prediction for the complementary case that assumes the primacy of NREM sleep for neural reorganization and information processing. This will be dramatically different and distinguishable from the predictions for REM sleep, hence providing a clear indication for when these processes occur during the sleep cycle.

### 1.3 Developmental changes in cerebral metabolic rate, synapses, and white matter

The theory for neural repair and reorganization developed above is fundamentally driven by the brain’s metabolic rate. In order to make the scaling relationships for sleep fully predictive, the only remaining unknown is the scaling exponent, *α*, that relates brain metabolic rate to brain size. Across species, the brain can be treated as a nearly autonomous organ with its own vascular system supplied primarily by a single carotid artery, much in the same way that the vascular system of the entire body is supplied through a single aorta. Following the theoretical derivation of the scaling relationship of metabolic rate for the whole body, this predicts 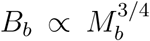, consistent with data for mature mammals (42, 68).

However, during early ontogeny the brain is undergoing rapid changes in size and synaptogenesis that require a relaxation of the power optimization and other constraints that determine how metabolic rate scales with size for mature organisms (69). Therefore, the canonical theory for the scaling of metabolic rate needs to be reformulated for the brain to recognize that ontogeny may not recapitulate phylogeny. Neural signaling and computation in the brain are extremely costly, accounting for 80 − 90% of its metabolic expenditure (70–72). These signals and computations are implemented through patterns of neuronal synaptic connectivity. It is therefore natural to focus on the number of synapses as a major driver of brain metabolic rate rather than the number of neurons (52). The primary function of these connections is to regulate electrical signals through axons that transmit information through neural networks to gather and process information, in order to learn and react (73). Crucially, as the number of synapses quickly increases in early development, they bring along associated increases in glial cells that are also highly metabolically active. Following this initial increase in the number of synapses, they are subsequently pruned away as part of the process of learning and reorganization in a way that is consistent with the Hebbian maxim that neurons that fire together, wire together (74).

Consequently, cerebral metabolic rate at early developmental stages is proportional to the total number of synapses already present plus the rate at which energy needs to be supplied to grow new ones. This is consistent with prior work showing the invariance of cerebral metabolic rate per synapse across development for mammals (72). Since most neurons in the adult brain are already present at or soon after birth, with only an extremely slow increase in their number during development and adulthood (75), the metabolic rate devoted to existing synapses at any given time is much greater than that needed to create new ones.

The increase in the number of synapses after birth largely represents the wiring together of pre-existing neurons, further emphasizing the dependence of changes in metabolic rate on synapse number rather than neuron number.

As a result, we predict that the metabolic rate of the brain should scale approximately linearly with the total number of synapses or, equivalently, that its mass-specific metabolic rate should scale linearly with synaptic density. Additionally, after birth the increase in brain mass derives largely from the increase in glial cell and neuronal spine mass within grey matter and through the myelination of axons within white matter (76). The primary function of glial cells is to support synaptic activity (77), so increases in white matter are driven by increasing synaptic demand. Analogously, increases in myelination reflect the need for increased speed and bandwidth of axonal information transfer as the number of synapses per axon increases. Hence, we expect synapse number to scale approximately linearly with white matter volume, *V*_*w*_:

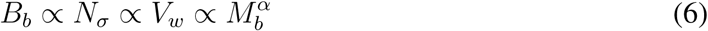

Intriguingly, previous studies across species have shown that the volume of white matter increases superlinearly (scaling exponent > 1) with that of gray matter, *V*_*g*_, across species (78). If a similar result holds during development, which we test below, this would predict superlinear scaling (*α* > 1) for brain metabolic rate with brain size. This result means that brain metabolic rate per gram of tissue or per cell is actually *increasing* during development, in marked contrast to all other scaling relationships between metabolic rate and brain or body size.

The reconciliation between such a superlinear scaling across development and a sublinear allometric scaling across species can be understood in two ways. First, in adults the number of connections scales linearly with the number of neurons across species, representing a roughly constant adult synaptic density that is realized after pruning is complete (79). Second, for adults the number of neurons in the brain scales non-linearly and approximately as the 3*/*4 power with brain size across species (80).

### 1.4 Dramatic phase transition in sleep function related to brain development

As discussed below, a major transition in brain development (81, 82) and growth occurs around 2 to 3 years old in humans that is associated with the stabilization of synaptic growth and connectivity (52, 79, 83–85). In our theory sleep is inextricably linked with brain development, function, and metabolic rate. Consequently, we predict a sharp and dramatic transition in the scaling of sleep function at about this age. Such a transition reflects a fundamental change in brain development that occurs shortly after the peak in synaptic density when connectivity patterns in the brain begin to stabilize and sleep function shifts from being dominated by neural reorganization towards neural repair and maintenance. Indeed, because the fraction of REM sleep is predicted to change with size in the regime when neural reorganization dominates but be invariant for neural repair, we might expect a classic phase transition, analogous to when water freezes to ice. Mathematically, this would reveal itself as a discontinuity in the first derivative in the fit. Below we present an analysis of the data to confirm this remarkable prediction and show that it occurs at about 2.4 years old. To our knowledge, this sharp transition in sleep function, its simultaneity with the transition in rates of synaptogenesis and synapse density, and the associated scaling behavior have neither been previously predicted nor documented. This is remarkable given that this shift likely signals a profound shift in the function of sleep and the behavior of sleep processes.

## 2 Methods

### 2.1 Empirical Data

To conduct empirical tests of the predictions of our models, we surveyed the literature for available data on sleep times, REM sleep times, brain size, brain metabolic rate, body size, body metabolic rate, and other relevant factors for humans during growth from birth to adulthood. Altogether, compiled data contain about 400 points, mostly corresponding to an age range of 0-15 years. The study of Galland et al. (44) contained 105 data points for sleep times of humans from ages 0 to 12 years. The study took data from multiple individuals and provided error bars on sleep times as the mean ± 1.96 standard deviations to approximate 95% confidence intervals. Further data (40 data points) were obtained from Dekaban et al. (86) for brain weight for 0 to 20 year olds. Because we do not consider the effects of gender differences, we combine these data by calculating the mean of the female and male brain weights and body weights. Data for the percentage of REM sleep time across ontogeny and before 18 years old were found in Roffwarg et al. (45). In addition, sleeping metabolic rate (SMR) values from 0 to 1 year old are taken from Reichman et al. (87). They performed repeated measurements of SMR at 1.5, 3, 6, 9 and 12 months of age in 43 healthy infants. To better test for connections between white matter, synaptic densities, and cerebral metabolic rate, we also use ontogenetic data for cerebral metabolic rate (28 data points) and synaptic density (12 data points) for 0 to 15 years old from Feinberg et al. (52), as well as data for white-matter and grey-matter volume (88) from 0 to about 3 years old. Because numerical values or tabular data were rarely published for these studies, the software DataThief was used to collect data from graphs. Moreover, when combining two different datasets, if the ages were not completely aligned, we used interpolation to obtain values at the same age.

### 2.2 Data Analysis

To illuminate patterns in these data, we test for relationships between sleep time, brain size, and metabolic rate in humans. More specifically, we analyze the data from these different sources by constructing plots, calculating correlations between variables, and measuring slopes and exponents to test if empirical values match our theoretical predictions.

We focus our analysis on ages 0 to 12 years old because the data show that the relevant variables mostly stabilize after 12 years old. In doing our analysis, we note that this period itself can be split into two distinct regimes and discuss how the relative roles of repair/clearance and reorganization shift during this time. By dividing the data into two separate regimes, the logarithmic plots closely follow a straight line for each of these two regimes. Because biological and physical changes are typically continuous, we require that the line before and the line after the transition connect to each other in a continuous fashion. We first choose this intersection point (*x*_0_, *y*_0_), and we then use two lines *y* = *k*_1_(*x* − *x*_0_) + *y*_0_ and *y* = *k*_2_(*x* − *x*_0_) + *y*_0_ to fit the data. We determine the best value of *k*_1_ and *k*_2_ as well as the intersection point (*x*_0_, *y*_0_) through a minimization of the sum of the squared errors (SSE) (see Supp. Info. 3).

## 3 Results

Brain metabolic rate is fundamental to our theories of sleep for neural repair and for neural reorganization. We thus begin by analyzing our collected dataset to determine the scaling relationships between brain metabolic rate, total number of synapses, volume of white matter, and brain size (see Eq. (6)). These will be used to test our predictions and determine the exponent, *α*, needed to complete the quantitative predictions for sleep times expressed in Eqs. (1-5). We divide the analysis of sleep data into two distinct sleep phases based on the predicted and statistically determined transition that occurs between 2 and 3 years of age. Fig. (1) shows three plots that evaluate our main predictions for these quantities in the early development phase prior to this transition.

**Figure 1:**
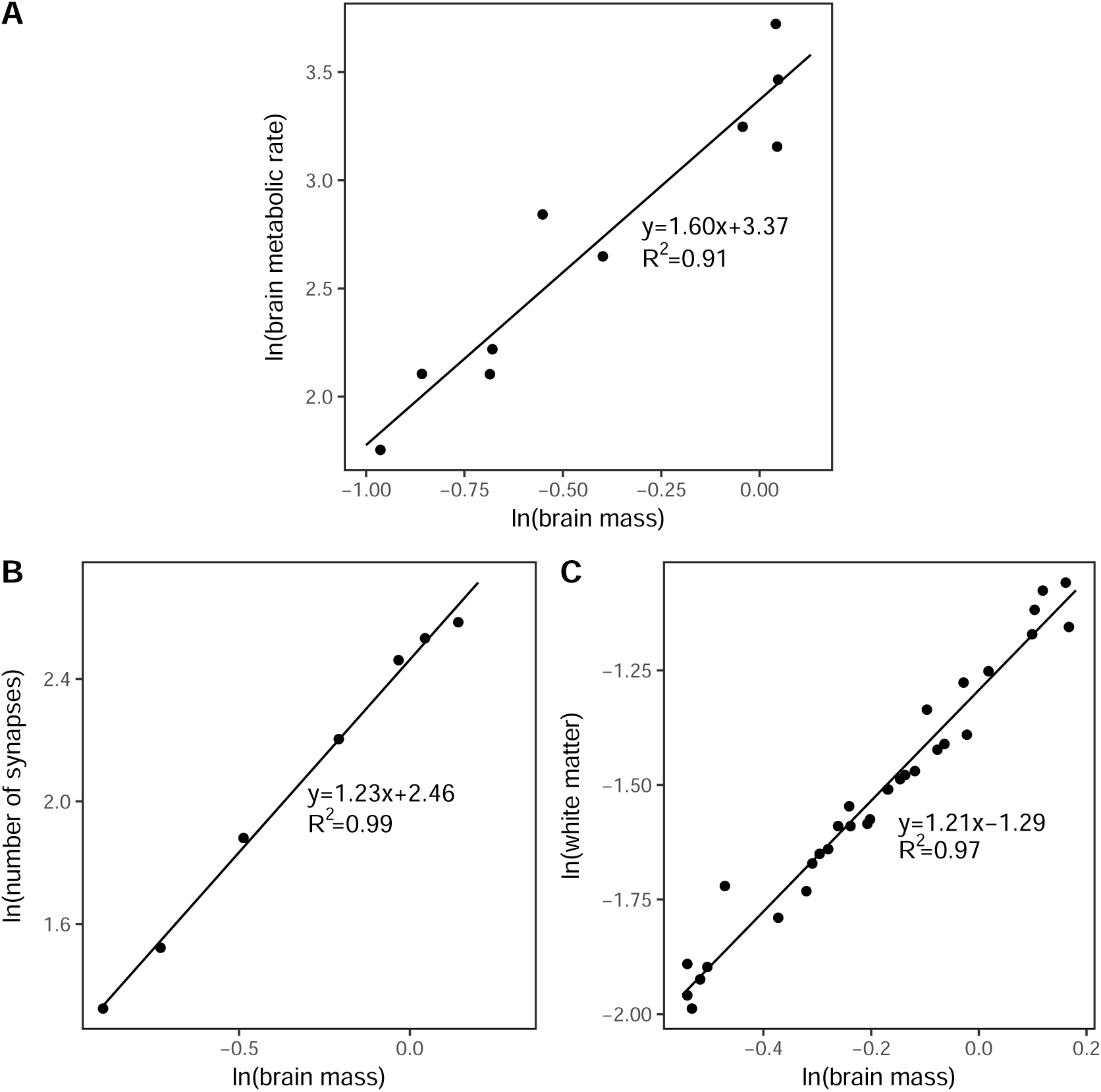
A. Plot of the logarithm of cerebral metabolic rate versus the logarithm of brain mass before transition with measured slope 1.60. B. Plot of the logarithm of number of synapses versus the logarithm of brain mass before transition with measured slope 1.23. C. Plot of the logarithm of white matter volume versus the logarithm of brain mass before transition with measured slope 1.21.

i. Fig. (1A) shows a logarithmic plot of the brain’s sleeping metabolic rate versus its mass. This reveals a remarkable superlinear behavior with an exponent, *α* = 1.60 ± 0.40, confirming our prediction of superlinear scaling based on the scaling of white and grey matter. Most notably, it runs strongly counter to all the normal patterns of allometric scaling relationships across species that are invariably sublinear (i.e., with exponents <1) (65–67, 89–91). This results from the brain becoming increasingly energetically more costly during development and stands in marked contrast to the energetics of all other tissues and organs in the body where economies of scale dominate. In that case, the larger the organism (or organ) the *less* metabolic power is required per unit mass of tissue. Superlinear scaling, on the other hand, means exactly the opposite: the larger the organism (or, in this case, the brain) the *more* metabolic power is required per unit mass of tissue or per cell.
ii. Fig. (1B) shows a similar logarithmic plot for the number of synapses versus brain mass. The number of synapses is simply the product of synaptic density, *ρ*_*σ*_ - usually measured with respect to a local section of grey matter volume - and the volume of grey matter in the brain, *V*_*g*_: *N*_*σ*_ = *ρ*_*σ*_*V*_*g*_. Fig. (1B and Supp. Info. 4) yields a scaling exponent of 1.23 ± 0.09, which is consistent with our prediction, Eq. (6), from the scaling of brain metabolic rate as well as with the scaling of white matter with grey matter across species.
iii. Lastly, we evaluate our predictions based on a much more comprehensively measured property, namely the volume of white matter as a function of brain mass. Fig. (1C) shows a plot for this relationship that reveals a scaling exponent of 1.21 ± 0.08, consistent with our predictions and the other two estimates of *α*.

These results show that the value of the superlinear exponent *α* is in the range from 1.20 to 1.60. We now use this in Eqs. (1-5) to predict how sleep time ratios, such as *t*_*S*_*/t*_*A*_ and *t*_*R*_*/t*_*S*_, scale with brain and body size. Recall that the predictions depend on whether sleep function is dominated by neural repair or neural reorganization. If it is based on neural repair, Eq. (1) predicts that *t*_*S*_*/t*_*A*_ scales with an exponent between 0.20 and 0.60, and that the fraction of REM sleep (*t*_*R*_*/t*_*S*_) is invariant with respect to brain mass. In contrast, if neural reorganization dominates, we predict from (4) that *t*_*R*_*/t*_*A*_ scales with an exponent between -1.20 and -0.40 if driven primarily by REM sleep, whereas if it is driven primarily by NREM sleep, *t*_*NR*_*/t*_*A*_ scales with an exponent between -1.20 and -0.40. This provides a remarkably clean test for discerning between different underlying mechanisms for sleep, and whether they occur during REM or NREM sleep.

In Fig. (2) we analyze data and provide strong statistical evidence for the existence and sharpness of the transition from early to late development. To identify the location of the transition and measure its sharpness, we focus on two independent measures of sleep that are related to total sleep time and the NREM/REM sleep trade-off, *t*_*S*_*/t*_*A*_ and *t*_*NR*_*/t*_*A*_. To determine the transition point in brain mass for each of these sleep ratios, we choose all possible break points in the data for *M*_*b*_ and calculate the corresponding Sum of Squared Errors (SSE) of the residuals from the two best fit straight lines on either side of each possible break point. As observed in Fig. (2), there are unique and sharp minima at almost exactly the same value of *M*_*b*_ in both plots, corresponding to the same age in development. Based on these results, we identify the transition point to be at 2.4 years old, consistent with the age range of 2 to 3 years old that corresponds to many known transitions in brain development (81–84, 92).

**Figure 2:**
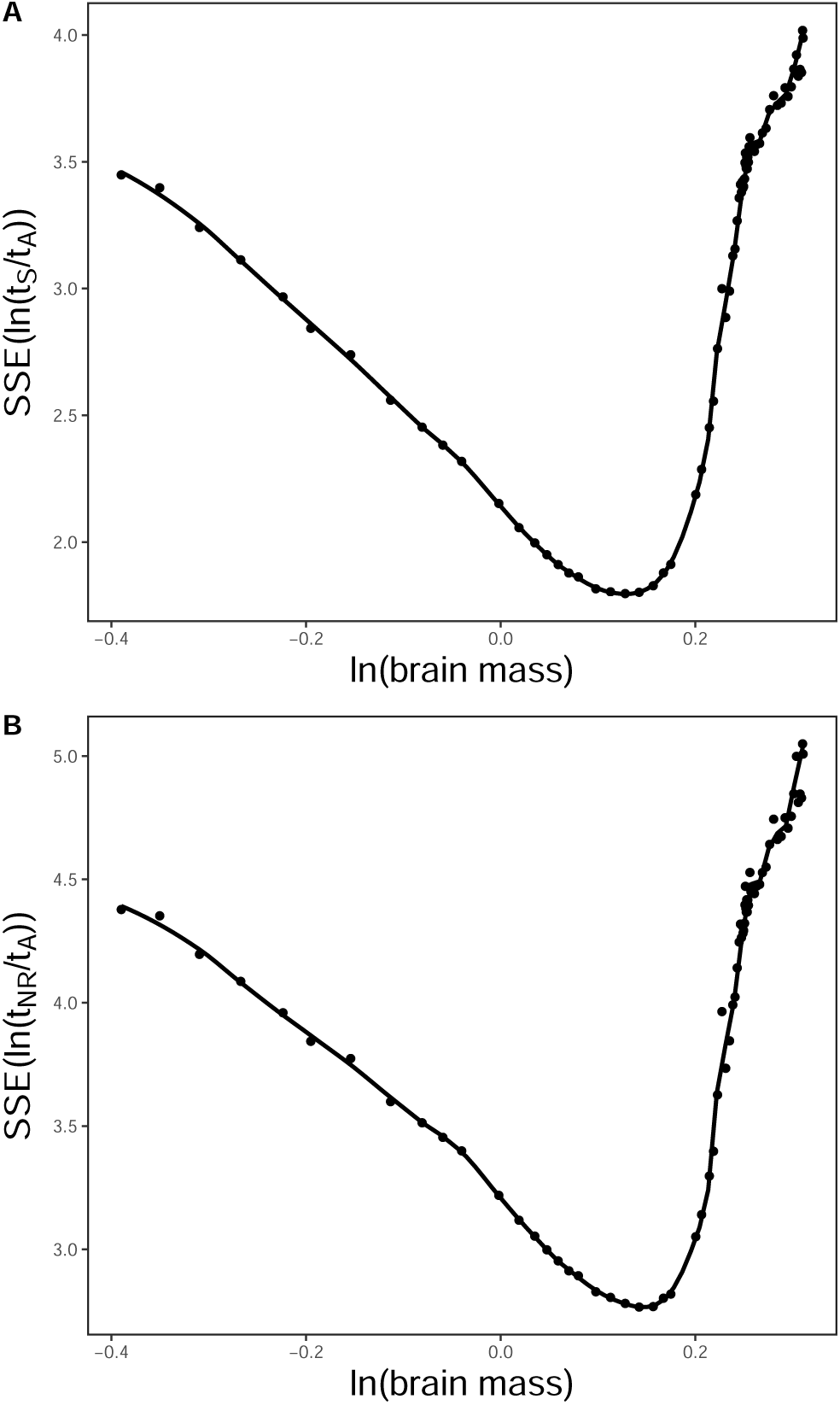
Plots of the Sum of Squared Errors (SSE) for the residuals of the two best-fit lines to data for (A) ln(*t*_*S*_ */t*_*A*_) and (B) ln(*t*_*NR*_*/t*_*A*_) on either side of a break point in the lines that corresponds to that value of the logarithm of brain mass (*M*_*b*_). The minimum of each curve is identified as the transition point that divides sleep function into early and late developmental stages as described by our theory. These minima are unique and have values of *M*_*b*_ = 1.14 kg for the transition in *t*_*S*_ */t*_*A*_ and *M*_*b*_ = 1.15 kg for the transition in *t*_*NR*_*/t*_*A*_, corresponding to ages of 2.4 to 2.5 years old respectively.

In Fig. (3) we present plots of the various sleep time ratios versus brain mass, demonstrating clearly that all of the data for sleep time ratios exhibit a clear transition from early to late development. Using our compiled developmental data for sleep in humans, we test predictions from the theory and are thereby able to determine the underlying mechanisms of sleep.

**Figure 3:**
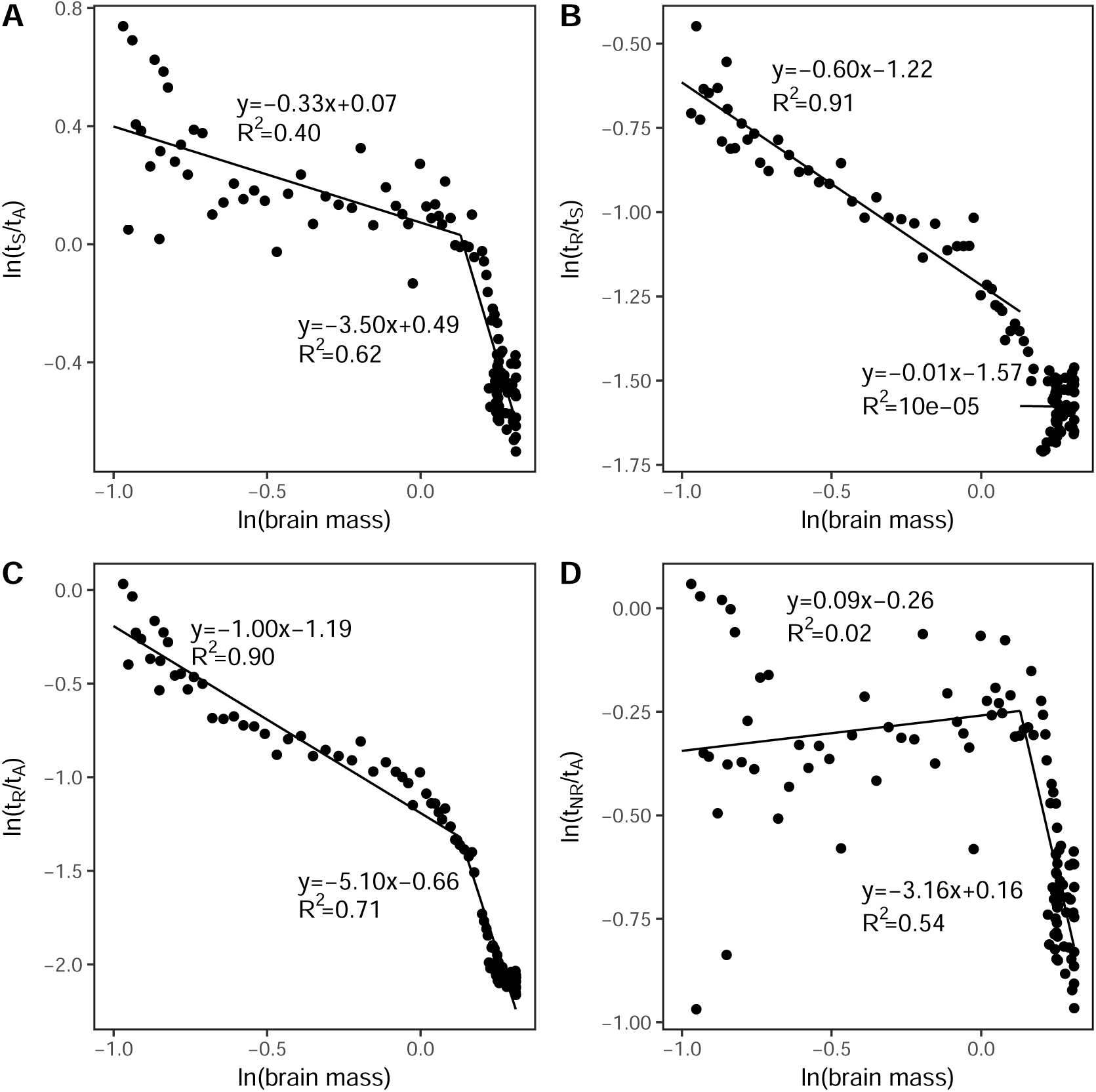
A. Plot of the logarithm of the ratio of total sleep time to total awake time per day versus the logarithm of brain mass with measured slope -0.33 before transition and -3.50 after. B. Plot of the logarithm of the ratio of REM sleep time to total sleep time per day versus the logarithm of brain mass with measured slope -0.60 before transition and -0.01 after. C. Plot of the logarithm of the ratio of REM sleep time to total awake time per day versus the logarithm of brain mass with measured slope -1.00 before transition and -5.10 after. D. Plot of the logarithm of the ratio of NREM sleep time to total awake time per day versus the logarithm of brain mass with measured slope 0.09 before transition and -3.16 after.

Fig. (3A) shows that the scaling exponent for the ratio of sleep to awake time, *t*_*S*_*/t*_*A*_, during early development (< 2.4 years) is −0.33 ± 0.07, in the opposite direction (decreasing with size rather than increasing) and strongly at odds with the predictions from neural repair. Similarly, in Fig. (3B) we see that the scaling exponent for the fraction of REM sleep, *t*_*R*_*/t*_*S*_, is −0.60 ± 0.06 during early development, again in complete contradiction to the invariance predicted from neural repair. On the other hand, from Fig. (3C) the ratio of REM sleep time to awake time, *t*_*R*_*/t*_*A*_, has an exponent of −1.00 ± 0.05, consistent with the prediction that assumes sleep function is primarily driven by neural reorganization during REM sleep. Finally, as a consistency check on this, Fig. (3D) reveals that the corresponding exponent for the ratio of NREM sleep time to awake time, *t*_*NR*_*/t*_*A*_, is 0.09 ± 0.09, consistent with it being an invariant and strongly counter to the predictions assuming that sleep function is primarily driven by neural reorganization during NREM sleep.

As a further test of our predictions, we return to Equation (5). Because the observed scaling of *t*_*S*_*/t*_*A*_ has an exponent of −0.33, the theory based on REM sleep being for neural reorganization would predict that the exponent for the fraction of REM sleep, *t*_*R*_*/t*_*S*_, should be between -0.87 and -0.07. This differs significantly from the invariance predicted from neural repair and implies that the empirically measured exponent of -0.60 provides additional support for sleep function during early development being tied to neural reorganization in REM sleep.

To summarize: when theoretical predictions are confronted with empirical data, the only consistent mechanism is that *sleep function throughout early development is primarily driven by neural reorganization during REM sleep*. Equally importantly, all other mechanisms are soundly rejected as can be seen by comparing measured scaling exponents and their confidence intervals from Figs. 1-3 with predictions from our theory (see Table 1).

**Table 1:** Early Development (<2.4 years) Summary of the key empirical results and theoretical tests of our model for the various ratios of sleep times in the first column during the period of early development (< 2.4 years old). The second column contains the values and 95% confidence intervals (CI) for the scaling exponents as determined from direct empirical data, whereas the third through fifth columns contain the ranges of predicted values for the scaling exponents based on theories that sleep function is primarily for neural reorganization in either REM (3rd column) or NREM (4th column) sleep or that it is primarily for neural repair (5th column). The range of predicted values is calculated in each case using the three best-fit estimates for the scaling exponent *α* from Fig. (1). NA denotes that the corresponding theory makes no prediction for that specific variable. Predictions that match data are in red. For these data, the predictions of the theory that sleep function during early development is primarily for neural reorganization in REM sleep are all supported, whereas the predictions are all rejected that during early development sleep function is either primarily for neural repair or for neural reorganization during NREM sleep.

| Ratio | measured exponent | REM reorg. prediction | NREM reorg. prediction | repair prediction |
| --- | --- | --- | --- | --- |
| $t_S/t_A$ | $-0.33 \pm 0.07$ | NA | NA | 0.20 to 0.60 |
| $t_R/t_S$ | $-0.60 \pm 0.06$ | **-0.87 to -0.07** | NA | 0 |
| $t_R/t_A$ | $-1.00 \pm 0.05$ | **-1.20 to -0.40** | NA | NA |
| $t_{NR}/t_A$ | $0.09 \pm 0.09$ | NA | -1.20 to -0.40 | NA |

The result that REM sleep time takes up about 50% of total sleep time for newborns, whereas people older than 50 years spend only about 14% − 15% of their sleep time in REM (45), is a particularly striking result. Indeed, this ontogenetic change is a fundamentally different pattern than that observed phylogenetically, i.e. across species, in which the fraction of time spent in REM sleep does not change from mice to whales. Yet the ontogenetic change is consistent across phylogeny (45, 93–95). The decline in the fraction of REM sleep strongly suggests the decreasing importance of reorganization as a function for sleep beyond about human age 2.4 years old, and correspondingly, the ascendance of repair and/or clearance as the primary function in later development (Table 2). That is, as we grow, the dominance of sleep by processes for neural reorganization transitions to the dominance by neural repair and clearance. To test this, we fit the data for *t*_*R*_*/t*_*S*_ after the transition point (Fig. (3B)) and find that it has a slope not significantly different from 0 and therefore consistent with it being an invariant as it is across adult mammals.

**Table 2:** Late Development (>.4 years) Summary of the key empirical results and theoretical tests of our model for the various ratios of sleep times in the first column during the period of late development (> 2.4 years old). The second column contains the values and 95% confidence intervals (CI) of the scaling exponents as determined from direct empirical data, whereas the third through fifth columns contain the ranges of predicted values for the scaling exponents based on theories that sleep function is primarily for neural reorganization in either REM (3rd column) or NREM (4th column) sleep or that it is primarily for neural repair (5th column). The 95% confidence intervals for the predictions are derived from the confidence intervals determined for the scaling exponent *α* = −1.70 ± 1.66 in later development (Fig. (S2)). NA denotes that the corresponding theory makes no prediction for that specific variable. Predictions that match data are in red. For these data, the predictions of the theory that sleep function during early development is primarily or neural repair and clearance are all supported, whereas the predictions that during early development sleep function is primarily for neural reorganization in REM sleep or NREM sleep are all rejected.

| Ratio | measured exponent | REM reorg. prediction | NREM reorg. prediction | repair prediction |
| --- | --- | --- | --- | --- |
| $t_S/t_A$ | $-3.50 \pm 0.23$ | NA | NA | **-2.70 ± 1.66** |
| $t_R/t_S$ | $-0.01 \pm 0.52$ | $8.90 \pm 3.55$ | NA | **0** |
| $t_R/t_A$ | $-5.10 \pm 0.20$ | $5.40 \pm 3.32$ | NA | NA |
| $t_{NR}/t_A$ | $-3.16 \pm 0.26$ | NA | $5.40 \pm 3.32$ | NA |

Moreover, if we try to fit a line through these data to connect it with the line at our transition point, we obtain an *R*^2^ value of -0.33 (the negative sign being due to the fixing of the *y*-intercept), indicating a terrible fit. Taken together, this means that in later development, as well as across species, the scaling of the fraction of REM sleep is consistent with the prediction of it being invariant based on the importance of neural repair and clearance for sleep function. Furthermore, the fits indicate there is an actual discontinuity in the slope (i.e., first derivative) of this property, corresponding in physics terminology to a true phase transition (96) and indicative of a seismic change in sleep and brain function at this early age of around 3 years old.

As further support for the idea that sleep function in later development is for neural repair and clearance, we measure the scaling exponent of brain metabolic rate, *α*, beyond this transition (Supp. Info. E) and find a value of −1.70 ± 1.66. Using this in Eq. (1) predicts that *t*_*S*_*/t*_*A*_ should scale in this regime with an exponent of −2.70 ± 1.66, which is consistent with the value of −3.50 ± 0.12 measured in Fig. (3A). Altogether, this provides compelling evidence in favor of sleep function being primarily for neural repair and clearance during later development, beyond about 3 years old, and also into adulthood and across species (1, 42).

In summary, our main results are to

1. identify the exact transition point in the function of sleep from reorganization to repair in the brain and recognize that it tightly corresponds to transitions in brain development.
2. quantitatively demonstrate that this transition, which occurs at 2.4 years old, is remarkably sharp and analogous to a phase transition, or tipping point, as when water freezes to ice.
3. show that evidence supports the REM-reorganization theory of sleep prior to this early transition and strongly exclude both NREM-based reorganization and repair-driven mechanisms.
4. show that theories for the function of sleep in late development that are based on neural reorganization during either REM or NREM sleep are strongly excluded by the data.

## 4 Discussion

Sleep is such an engrained and necessary part of our lives that we often take its functions and origins for granted. Presuming that sleep evolved to serve some primary function, it is almost certain that multiple physiological functions have piggy-backed onto this pervasive and time-consuming feature of animal life. Here, by deriving a novel theory, compiling comprehensive data on sleep and brain development, and quantitatively comparing sleep ontogeny with sleep phylogeny, we illuminate the dominant functions of sleep and how they change through development. Infants spend a much greater percentage of time in REM sleep compared with older children and adults. This finding suggests that REM sleep is likely crucial for the initial growth of babies, and perhaps especially for the regulation of synaptic weights throughout the nervous system (97). These substantial changes in percent REM sleep across human growth are in stark contrast with the constant percentage of REM sleep observed across an enormous range in brain and body size for adult mammals (42). The large change in percent REM sleep across development is thus a key indicator that the function of sleep, and particularly of REM sleep, is very different during development than in adults. Indeed, it shows that ontogeny does *not* recapitulate phylogeny because ontogeny does not show qualitatively similar patterns to phylogeny for REM sleep. Rather, it differs from it in the most fundamental of ways (e.g., invariance versus rapidly changing) and exhibits a phase transition between early and late development.

In our analysis we divide development into two regimes: an early period of high plasticity accompanied by ongoing synaptogenesis and increasing myelination followed by a later period of declining plasticity, slow synaptic pruning, and increasing white matter integrity and stabilizing connectivity. Our new theory, mathematical models, and data analysis provide compelling evidence that these fundamental differences arise because sleep is used primarily for neural reorganization until about 2 to 3 years of age, at which point there is a critical transition and the function shifts sharply towards sleep being for repair and clearance. We identify the specific turning point as occurring at a surprisingly precise age of around 2.4 years old, reflecting a critical physiological or cerebral developmental change. In all cases, we see a sharp shift in the scaling of sleep during this period of early development that to our knowledge has never been conceptually or quantitatively connected to a shift in sleep function.

When looking at functional brain development in humans (85), Johnson found the first two years of life is the period that most of the pronounced advances in brain structure and behavior occur. Brains develop extremely dynamically in the first two years (84), and most brain structures have the overall appearance of adults by the age of around two. One notable exception is the delayed development of the prefrontal cortex, the onset of which perhaps corresponds with a surge in REM sleep around later puberty, which would be predicted by our theory. Overall brain size increases dramatically during the first two years of life and reaches 80 − 90% of adult size by the age of 2. All of the main fiber tracts are observed by 3 years old (85), and in frontal brain regions white matter changes most rapidly during the first two years. White matter is associated with cognitive function (98), so the rapid change of white matter by the age of two helps to partly explain why reorganization might dominate before 2.4 years of age and then transition to a different stage. Other critical periods and transitions in early development with regard to learning and brain function are well known and of great interest, including the acquisition of language (81–84, 92, 99).

Our ontogenetic findings differ markedly from previous phylogenetic findings both in terms of the magnitude and sometimes the direction of changes and the corresponding scaling exponents. These results are quite surprising, yet by transitioning from our model for neural reorganization to one for repair and clearance, we are able to simultaneously explain the scaling of sleep time across species as well as across growth. Indeed, repair/clearance (30–32, 100, 101) and reorganization both occur throughout growth, and in analyzing data and building our theory, we hypothesized and showed how each of these dominates during specific developmental stages: reorganization dominates at early ages whereas repair and clearance dominate at later stages. Remarkably, our theory explains the scaling with brain and body mass in these two different regions for quite different reasons. For neural reorganziation, the scaling arises due to the mismatch between the sub-linear scaling of whole body metabolic rate, which drives information transfer to the brain, and the super-linear scaling of synapse number and white matter volume. In contrast, for repair and clearance, the scaling arises because the repair and clearance mechanisms are proportional to brain mass, while the damage rate is proportional to brain metabolic rate that scales nonlinearly with brain mass, creating a different type of scaling relationship. These multiple origins of the scaling of sleep properties during different periods of life history, and how they arise from different sleep functions, is crucial because it allows us to match our different theories to the proposed functions for sleep. Another crucial difference is that the scaling exponents manifest as a steep, super-linear increase in brain metabolic rate with brain size during early development followed by a subsequent decline in later development.

Given the relative simplicity of the theory, the various sleep and cerebral properties predicted by our model match empirical data surprisingly well. We are not yet able to predict all measured sleep properties, but our agreement for such a diverse set of characteristics during ontogenetic development as well as across phylogeny in adults is impressive. This lends credence to our assumptions and to the quantitative, mathematical framework that we developed.

One of our most compelling findings is not just that there is a transition, but how sharp that transition is, leading to complete reversals in direction for scaling relationships and also percent REM now suddenly changing instead of being invariant as it is across species. Although sleep always involves a loss of consciousness and characteristic electrical activity, our results suggest that the underlying dynamics of sleep may change fundamentally around 2 to 3 years of age. During early development, when substantial synaptogenesis is occurring, connections between neurons are likely transitioning from more short-range correlations (e.g., spatially localized circuits or networks) to more long-range connections (e.g., whole-brain) (102, 103). Moreover, connections are much more plastic in early development, while they are much more solidified in later development. From this perspective, the brain is in a more fluid state at birth and “cools off” during early ontogeny until a critical point is reached at 2 to 3 years of age that corresponds to a more crystallized state of brain structure and dynamics.

A central feature of our approach is that it is quantitative, computational, predictive, and can be readily tested with empirical data. Our findings point towards new and exciting questions that require more studies. An open question is whether the same ontogenetic patterns in sleep exist for other species. Humans are known to be unusual in the amount of brain development that occurs after birth (75). Therefore, it is conceivable that the phase transition described here for humans may occur earlier in other species, possibly even before birth. Indeed, fetuses sleep a very large amount of the time (104), but it may be exceedingly difficult to take precise measurements of metabolic rate or brain mass and thus observe this shift in other species before birth. Measurements for growth and development in rats, zebra finches, drosophila, c. elegans, and many other species (27, 105, 105–107) are needed to test how well our theory generalizes to development in other species and the extent to which these shifts really are phase transitions.

## Supporting Information (SI)

### 4.1 Equation for mass dependence of total sleep time, *t*_*S*_

We can solve Eq. (1) in the main text by substituting *t*_*S*_ = 24 hours − *t*_*A*_ to obtain

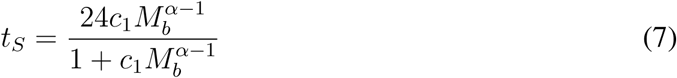

where 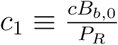 with parameters as defined in the main text and *B*_*b*_,_0_ representing the normalization constant for relating brain metabolic rate, *B*_*b*_ to brain mass, *M*_*b*_. Across species, the scaling exponent is sublinear, *α* < 1, so *α* − 1 < 0, resulting in *t*_*S*_ ∼ 24hours when *M*_*b*_ becomes increasingly smaller and 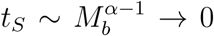 when *M*_*b*_ becomes increasingly larger. In strong contrast, within species, the scaling exponent is superlinear, *α* > 1, so *α* − 1 > 0, resulting in *t*_*S*_ ∼ 24hours when *M*_*b*_ becomes increasingly larger and *t*_*S*_ 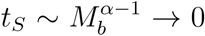 when *M*_*b*_ becomes increasingly larger. These limits make it strikingly clear that these predictions are the opposite of real patterns of sleep during growth.

### 4.2 Scaling of brain mass with body mass during early development

We note that data confirm that the brain mass of infants scales with their body mass as a power law whose exponent is 0.72, as shown in Fig. (S1), and that this value is very close to the value of 3*/*4 that was used in our predictions for the scaling of sleep properties.

### 4.3 Determination of transition age

Surveying the plots of our data, we note the sharpest and most dramatic transition of any of the sleep variables is for the ratio of NREM sleep time to awake time or for REM sleep time to awake time, strongly supportive of a transition in sleep function. Indeed, this ratio remains unchanged as a function of brain mass during the earliest stages of development until a transition occurs after which it steeply decreases. Thus, beyond this transition the fraction of REM sleep sharply decrease as a function of brain mass and age. Fig. (2) in the main text displays the data for this relationship and the dramatic transition from one scaling regime to another is clearly evident. Our fitting procedure identifies the transition age as 2.4 years old. Notably, there is a similar transition at roughly the same age in the ratio of sleep time to awake time (Fig. (3A) in main text) and of REM sleep time to awake time (Fig. (3C) in main text). Because the ratio of sleep to awake time and REM to awake time are primary drivers of sleep relationships within our models, these findings are consistent with a general shift in all sleep properties at a common age. For this reason, we use the following equations to determine the transition point.

**Figure S1:**
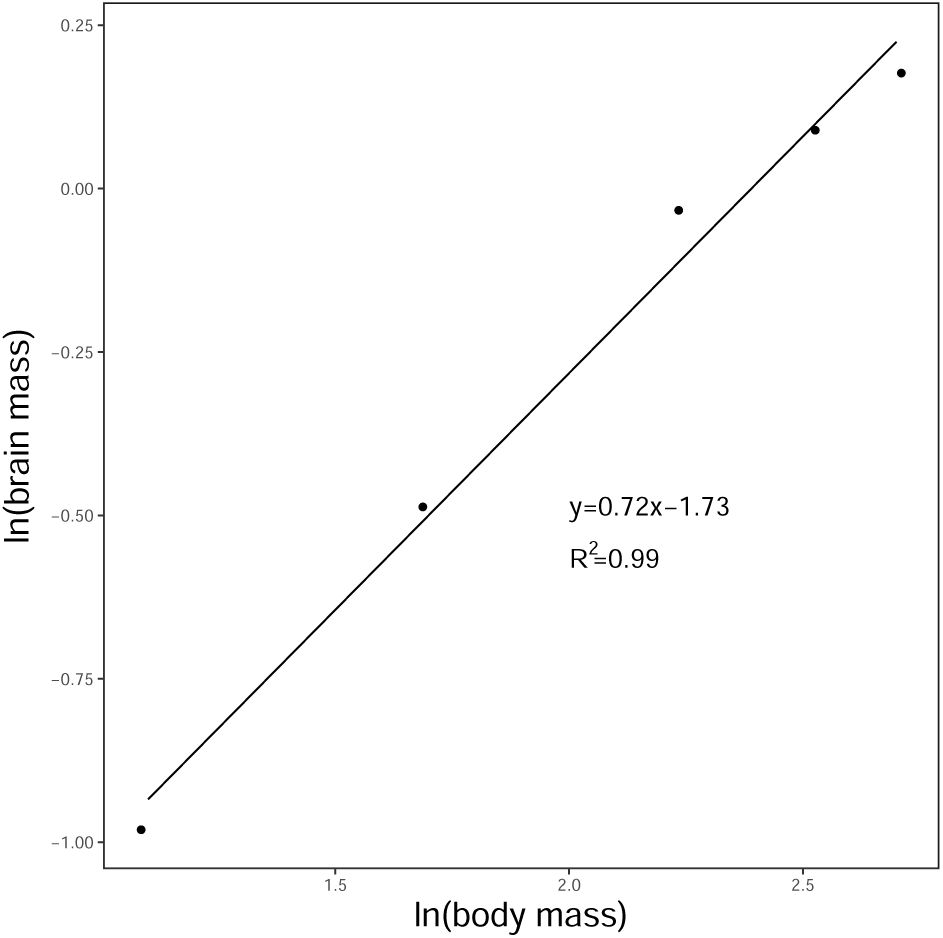
Plot of the logarithm of brain mass versus the logarithm of body mass before the transition with the measured slope 0.72.

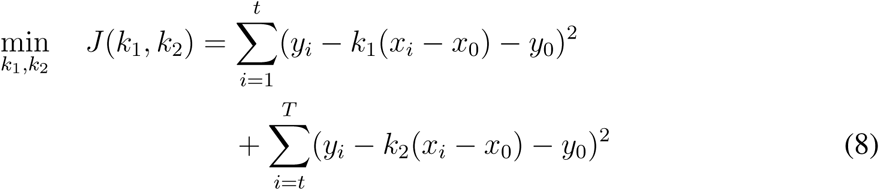

Taking the derivative of *J*, we have

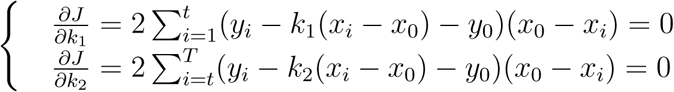

which implies that the optimal *k*_1_ and *k*_2_ with the corresponding transition point (*x*_0_, *y*_0_) are

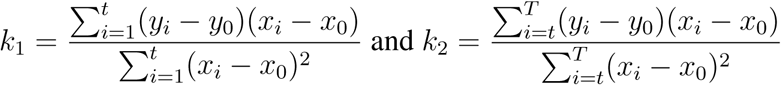

### 4.4 Raw data for scaling of number of synapses in early development

Our plots for number of synapses, *N*_*σ*_, in the main text were calculated from raw data for synaptic density, *ρ*_*σ*_, and gray matter volume, *V*_*g*_. Plots of the raw data for these two parameters are shown in Fig. (S2). Based on the scaling exponents of 0.39 and 0.91 measured in these plots, we would predict that 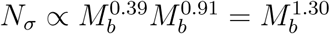, which is consistent with the measured scaling exponent for *N*_*σ*_ of 1.23 ± 0.09 from Fig. 1B in the main text.

**Figure S2:**
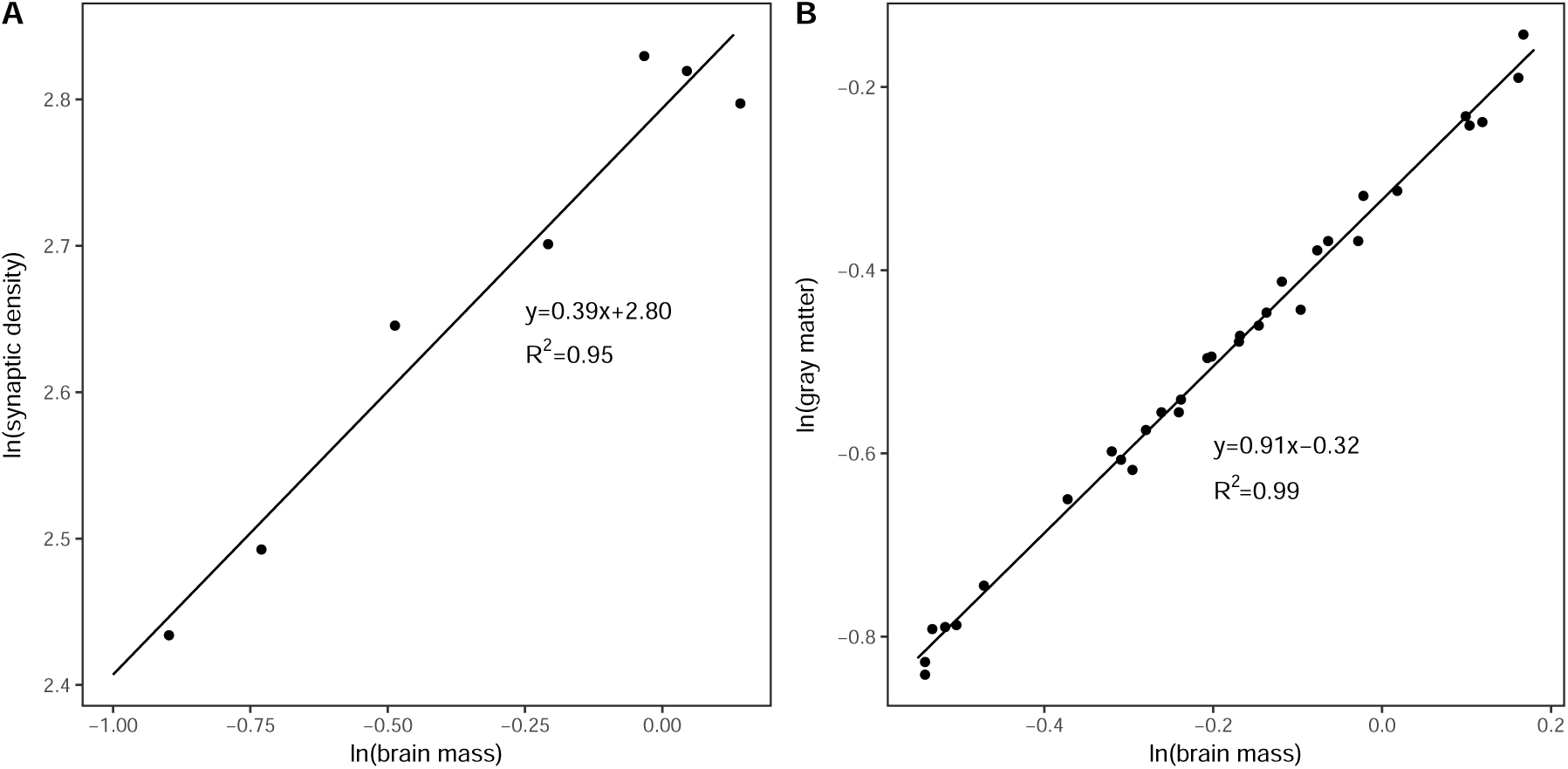
A. Plot of the logarithm of synaptic density versus the logarithm of brain mass before transition with measured slope 0.39. B. Plot of the logarithm of gray matter versus the logarithm of brain mass before transition with measure slope 0.91.

### 4.5 Scaling of brain metabolic rate during later development

Below we show a plot and measured scaling exponent for how brain metabolic rate, *B*_*b*_, changes with brain size, *M*_*b*_. This scaling exponent as an additional check of our theory for sleep being for neural repair during later development. We caution against much certainty or interpretation about the exact value (−2.70) of the exponent and its being significantly larger than the classic value of −0.25. Importantly, it should be noted that the range in brain mass over this time period of development is extremely small and the data are quite limited. Both of these properties of the data call into question the use of logarithmic variables to determine the exact magnitude of the scaling exponent in this case (91, 108), and the large 95% CI indicate the high degree of uncertainty in the exact value of this estimate.

**Figure S3:**
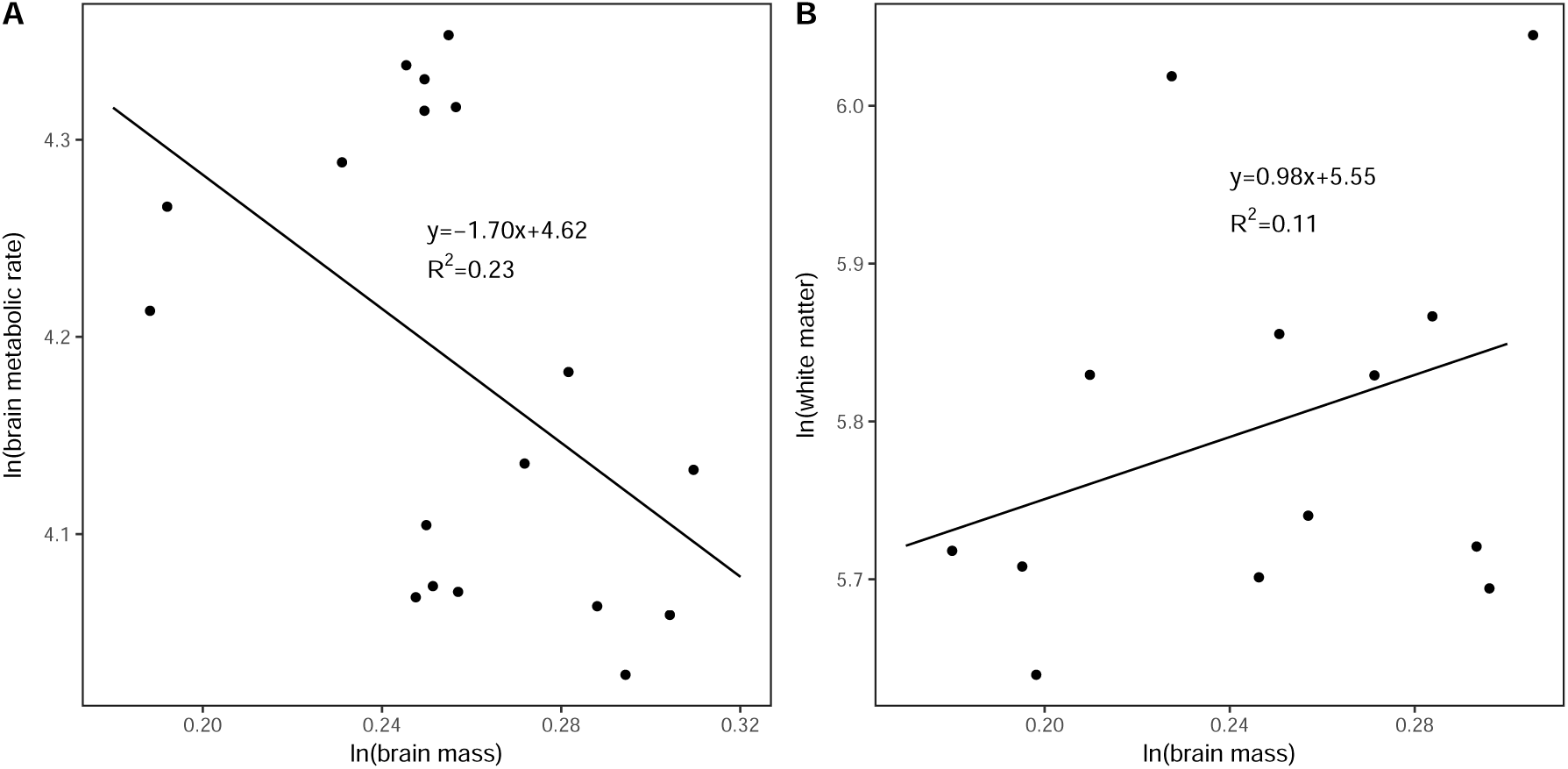
A. Plot of the logarithm of brain metabolic rate versus the logarithm of brain mass after 2.4 years old with the measured slope -1.70. B. Plot of the logarithm of the white matter volume versus the logarithm of brain mass after 2.4 years old with the measured slope 0.98.

**Figure S4:**
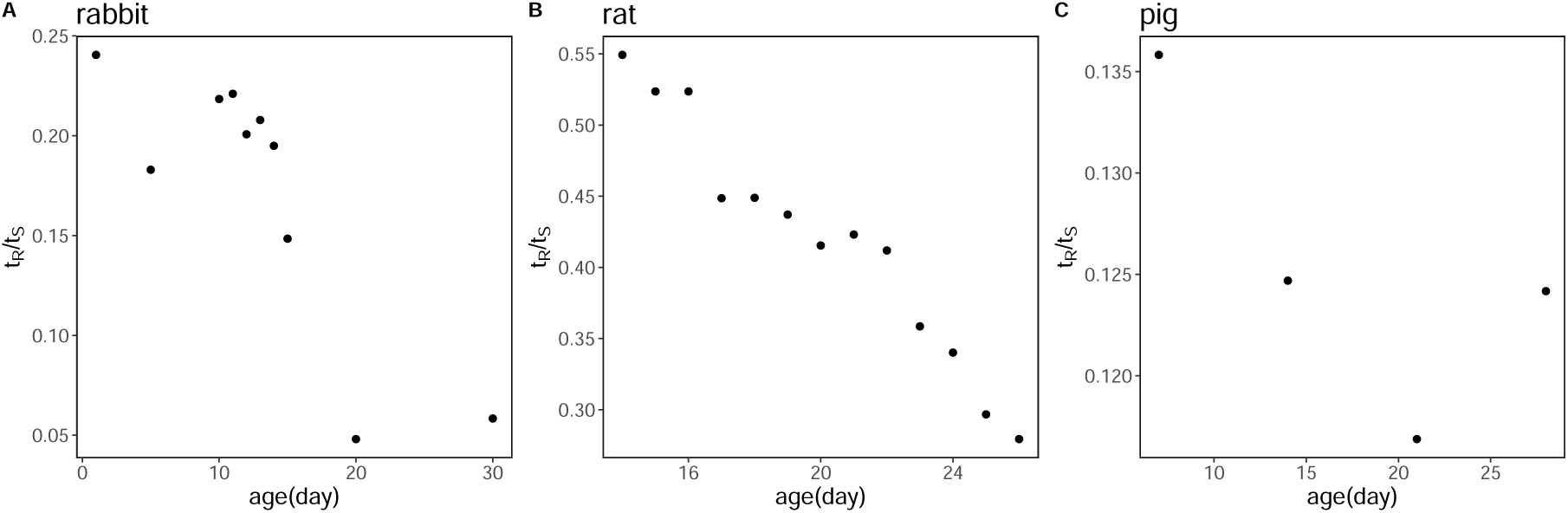
Plots of the ratio of REM sleep time to total sleep time per day versus the age for rabbit, rat, and pig, respectively.

### 4.6 Scaling of sleep across development in other species

We collect sleep data for three other mammals: rabbit (109), rat (110), and guinea pig (111). Fig S4 is a plot of the ratio of REM sleep time to total sleep time per day versus the age (days) for these three species.

## Acknowledgments

VMS acknowledges funding from an NSF DBI CAREER Award (1254159), and GBW would like to thank the National Science Foundation under the grant PHY1838420, the Eugene and Clare Thaw Charitable Trust, and Toby Shannan and CAF Canada for their generous support.

